# Mimicin, an antimicrobial protein encoded by mimivirus

**DOI:** 10.64898/2026.09.16.751764

**Authors:** Juliana Miranda Tatara, Hymonti Dey, Nisha Goyal, Ana Karoline Nunes Alves, Lara Ambrosio Leal Dutra, Abdeali M. Jivaji, Hans-Matti Blencke, Tor Haug, Frank O. Aylward, Jonatas Santos Abrahao, Matthias Horn, Gabriel Magno de Freitas Almeida

## Abstract

Antimicrobial peptides (AMPs) are innate defense molecules found in all domains of life. Giant viruses of amoeba are known to thrive among complex microbial relationships within its hosts cells, hinting at the existence of virus-derived antimicrobial strategies. Here we show that viruses belonging to the *Mimiviridae* and *Marseilleviridae* families contain a higher density of *in silico* predicted AMPs per genome size than other viruses of amoeba. The investigation of potential AMPs led to the description of Mimicin, a taxonomically restricted 74 amino acid long protein coded by few mimiviruses. Mimicin contains three smaller predicted AMP sequences within it and has a broad *in vitro* antimicrobial activity against different bacteria, a yeast and two non-enveloped phages. In contrast, it has no activity against a marseillevirus, a mimivirus or human cell lines. When tested against bacterial endosymbionts co-cultured with *Acanthamoeba terricola*, Mimicin and its SIM-31 portion were able to control the attenuated *Protochlamydia amoebophila* but not the highly virulent *Parachlamydia acanthamoebae*. Based on deposited transcriptomic data, Mimicin is coded by an early gene more active during the beginning of the infection process. No structure could be predicted using Alphafold, while additional structural analysis indicate that Mimicin could be a highly disordered protein. In conclusion, we describe evidence of a biologically relevant antimicrobial activity derived from a giant virus. Mimicin highlights the relevance of AMPs from giant viruses for microbial ecology and opens the way for investigating their biotechnological and clinical potential.

## Introduction

Unicellular eukaryotic organisms are frequently engaged in symbiotic relationships with prokaryotes, viruses and other unicellular eukaryotes (**Karnkowska et al 2026**). Amoebozoans in particular are known to be prone to harbour endosymbionts, with frequent co-occurrence of microbes and giant viruses detected at the single-cell level (**Schulz et al 2025**). One possible explanation for the diverse microbial communities within amoeba cells is their active grazing through phagocytosis, a process used for feeding on microbes and exploited by some giant viruses as an entry mechanism (**Souza et al 2021, Queiroz et al 2022**). The diverse microbial communities comprised of amoeba and their intracellular microbes make them “melting pots” for microbial interactions (**Boyer et al 2009**) and likely source of evolutionary pressure for the evolution of synergistic and antagonistic strategies between its components.

There is evidence of conflict between giant viruses, virophages and bacteria within amoeba cells. It is well documented that co-infections of virophages and mimiviruses results in loss of fitness of the later (**Gaia et al 2013**). In tripartite cultures of *Acanthamoeba*, mimivirus and the intracellular bacteria *Babela massiliensis*, subculturing results in the loss of the bacterium. However, when the virophage Sputnik 1 is added to the system, mimivirus loses fitness and *B. massiliensis* thrives (**Slimani et al 2013**). Conversely, the bacterial symbiont *Parachlamydia acanthamoebae* (PAVD) blocks the replication of mimivirus, Viennavirus, or Tupanvirus in *Acanthamoeba* during coinfections (**Arthofer et al 2022**). Although no direct evidence of antibacterial-derived strategies from giant viruses is known, the subculturing of mimivirus for 150 passages in axenic conditions led to a genomic reduction of around 16%, indicating the loss of non-essential genes that may be linked to control of competitors within the amoeba cells (**Boyer et al 2011**).

Antimicrobial peptides (AMPs) are innate defense molecules found in the animal and plant kingdoms (**Zasloff 2002**), as well as in microbes. Microbial-derived AMPs are exemplified by archaeasins in archaea (**Torres et al 2025**), bacteriocins in bacteria (**Sugrue et al 2024**), peptaibols in fungi (**Szekeres et al 2005**) and amoebapores in entamoeba (**Leippe 1999**). The presence of AMPs in all domains of cellular life supports the idea that these molecules are widespread and important for innate antimicrobial defence. AMPs also occur in viruses. The T7-like coliphage phiKT codes for gp28, a disruptin that acts on bacterial outer membranes (**Holt et al 2022**). Peptides derived from the membrane-active domains of HIV gp41 and gp120 proteins have been shown to possess antimicrobial activity (**Cole et al 2003**). In 2014, data retrieval from protein databases to build up an AMP database identified a few putative viral AMPs from different phages and from hepatitis B virus (**Waghu et al 2014**). AMP and cell-penetrating peptide (CPP) prediction using structural viral proteins as sources revealed 426 potential AMPs and over 2000 CPPs (**Freire et al 2015**), with a CPP from Torque teno douroucouli virus and an AMP from rotavirus being the most active ones in antibacterial assays (**Dias et al 2017**). As far as we know, no AMP has been yet predicted from or experimentally found in giant viruses of protists.

Here we describe Mimicin, a small uncharacterized protein present in some mimiviruses that contains three predicted AMP sequences within it and possess antibacterial and differential virucidal activities. Mimicin is able to control the bacterial symbiont *Protochlamydia* but not the antiviral *Parachlamydia* symbiont in co-cultures with *Acanthamoeba*. Mimicin was found from a large-scale *in silico* screening of AMPs across all publicly available giant virus predicted proteomes, evidencing that AMP candidates among giant viruses are enriched in mimiviruses and marseilleviruses. Our description of an active antimicrobial protein from a giant virus opens the way for the characterization of other potential candidates within and beyond the *Mimiviridae*.

## Material and methods

### Prediction and synthesis of antimicrobial peptides from viral origin

All publicly available giant viruses’ proteomes from UniProt (available until January 2025) were downloaded as FASTA files and used for AMP prediction through PyAMPA (**Ramos-Llorens et al 2024**). The output of predicted AMPs was divided into three main categories (*Imitervirales, Algavirales* or *Pimascovirales*) and the following cut-offs were applied: peptide size between 10-25 amino acids, no cysteine residues within the sequence, hemolytic probability <0.1, toxicity probability <0.1 and cell-penetrating probability >0.50. Considering these cut-offs and trying to include peptides from different viral families, 15 peptides were selected and synthesized (DGpeptides, China). Three exceptions from the cut-offs used above were also synthesized. The first was one protein of 74 amino acids, named Mimicin, which contains three predicted AMP sequences within it. The first AMP within Mimicin was found with the cut-offs stated above (MRP-17) while the second (SIM-31) and third (VTG-17) did not fall within the thresholds but were also synthesized. The purity of all synthesized molecules was verified by high-performance liquid chromatography (HPLC). Sequences and other details of all synthesized peptides are indicated at the **Supplementary table 1**. All 18 peptides were diluted to 10 mg/ml in sterile Milli-Q water (Millipore, Burlington, MA, USA) before use.

### Antibacterial assays

All peptides were screened for antimicrobial activity against a panel of four Gram-positive and four Gram-negative bacteria, plus one yeast. Bacterial strains used were *Bacillus subtilis* 168 (ATCC 23857), *Escherichia coli* (ATCC 25922), *Klebsiella pneumoniae* (ST876), *Pseudomonas aeruginosa* (ATCC 27853), *P. aeruginosa* PA01 (ATCC 15692), *Staphylococcus aureus* (ATCC 9144), *S. borealis* (Hus23) and *S. epidermidis* RP62A (ATCC 35984). Antifungal activity was tested against the yeast *Candida albicans* (ATCC 10231). Minimal inhibitory concentration (MIC) values were determined using a modified broth microdilution assay based on the CLSI M07-A9 protocol. Overnight bacterial cultures were grown in Mueller-Hinton (MH) broth (Difco Laboratories, USA) and sub-cultured for 1–2 hours prior to use. The bacterial inoculum was adjusted to approximately 10^4^ CFU/mL in MH medium. Inoculum were added in a 1:1 ratio to 96-well plates (Nunc, Roskilde, Denmark) preloaded with two-fold serial dilutions of each peptide (final concentration range: 128–0.125 µg/mL). Plates were sealed with a breathable membrane (Breathe-Easy, Diversified Biotech, USA) and incubated in an EnVision 2103 microplate reader (PerkinElmer, Llantrisant, UK) at 35 °C, with OD_595_ recorded every 30 minutes for 24 hours following agitation. A similar approach was used to verify antifungal activity against *C. albicans*, but in this case the yeast was grown overnight in potato dextrose broth (Difco) supplemented with 2% D (+)-glucose (Merck, Darmstadt, Germany) at 25–30 °C with shaking at 200 rpm and diluted to approximately 4 × 10^5^ cells/mL prior to use.

### Mechanism of action studies

Nine *B. subtilis* 168 based biosensor strains were used to evaluate the mechanism of action of the peptides. Of these, eight strains carry chromosomally integrated promoter–bacterial luciferase operon fusions, each designed to report a specific global bacterial stress response: DNA replication (yorB), transcription (held), translation (yheI), cell walls and membranes (ypuA and liaI), fatty acid synthesis (fabHB), folic acid synthesis (panB) and viability (laiG) (**Virta et al 1995; Dey et al 2022; Juskewitz et al 2022**). Membrane integrity was assessed separately using a *B. subtilis* 168-based strain carrying the reporter plasmid pCSS962, which constitutively expresses a eukaryotic luciferase. Overnight cultures of the biosensor strains were prepared in MH medium supplemented with 5 µg/mL chloramphenicol (Merck KGaA, Darmstadt, Germany) and grown at room temperature. The cultures were subsequently used to inoculate fresh medium without antibiotics and grown until an OD600 of 0.1 was reached. For the membrane-integrity assay, aliquots of 90 µL of bacterial inoculum premixed with 1 mM D-luciferin potassium salt (Synchem Inc., Elk Grove Village, IL, USA) were added to black round-bottom 96-well microtiter plates (Nunc, Roskilde, Denmark) containing two-fold serial dilutions of the peptides (10 µL per well), resulting in final peptide concentrations ranging from 100 to 1.56 µg/mL. Luminescence resulting from bacterial membrane disruption was monitored every 18 seconds for 3 minutes in a Synergy H1 Hybrid Reader (BioTek, Winooski, VT, USA) for the membrane integrity assays or during 8 hours for the other biosensor strains. Chlorhexidine acetate (CHX; Fresenius Kabi, Halden, Norway) and Milli-Q water were used as positive and negative controls, respectively.

### Activity against bacterial symbionts

For testing the effect of Mimicin and SIM-31 against bacterial symbionts in the presence of amoeba cells, approximately 1 × 10^5^ *Acanthamoeba terricola* (formerly *A. castellanii* NEFF; ATCC 50373) trophozoites were seeded per well in 24-well plates and allowed to adhere in PYG medium (2 % proteose peptone, 0.1 M glucose, 0.2% yeast extract, 3.4 mM trisodium citrate dihydrate, 4 mM MgSo_4_.7H_2_O, 1.32 mM Na_2_HPO_4_.2H_2_O, 2.5 mM KH_2_PO_4_, 0.05 mM Fe (NH_4_)_2_(SO_4_)_2_.6H_2_O; pH 6.5) at 20°C for 2 hours. In parallel, Mimicin or SIM-31 at 32 µg/mL final concentration were preincubated with *Protochlamydia amoebophila* UWE25 (ATCC PRA-7) (**Collingro et al 2005**) or *Parachlamydia acanthamoebae* PAVD endosymbionts (**Arthofer et al 2022**) at MOI 20 in 100 µL PAS (2.1 mM NaCl, 0.016 mM MgSo_4_.7H_2_O, 0.027 mM CaCl_2_.2H_2_O, 0.8 mM Na_2_HPO_4_.2H_2_O, 1 mM KH_2_PO_4_) at 20°C for 30 minutes. The peptide-symbiont mixture was then added to a well seeded with amoebae in ~900 µL PYG. Three preincubation conditions were tested: Mimicin, SIM-31 and Milli-Q water as negative control. To assess amoeba toxicity in the context of these experiments, Mimicin or SIM-31 were preincubated in PAS alone at 20°C for 30 minutes and then added directly to amoebae in a separate 24-well plate. Amoeba cells were exposed to Mimicin or SIM-31 for 30 minutes, after which the peptides were removed by replacing the medium. For symbiont and virus co-infection experiments, cultures were simultaneously infected with mimivirus at MOI 10 (estimated by tissue culture infectious dose 50, TCID50) together with Mimicin or SIM-31 and *Parachlamydia*. Infections were synchronized by centrifugation at 1,000 *g* for 20 minutes at 20°C. After 10 minutes, cells were washed once with PYG and cultured in 1 mL PYG at 20°C for the indicated times post-infection. All conditions were set up in triplicates, and each experiment was independently repeated twice. At the indicated times, cultures were harvested to quantify mimivirus and symbiont DNA in the cellular fraction by dPCR and to determine amoeba counts. An aliquot of 10 µl was used to measure amoeba cell numbers (cells/mL) with a LUNA automated cell counter (LUNA FX_7_™ BioCat GmbH, logos, Germany). Cultures were then centrifuged to pellet cells: 10,000 × *g* for 10 minutes at 4°C for single symbiont infection, or 8,000 *g* for 30 minutes at 4°C for co-infections. Genomic DNA was extracted from the pellets using the QIAGEN Blood & Tissue Kit® according to the manufacturer’s instructions, eluted in 100 µL of elution buffer and stored at −20°C until dPCR analysis.

### Digital PCR (dPCR)

Mimivirus genome copies were quantified using 0.25 µM each of major capsid protein (MCP) targeting primers: MCP_ApolyphagaMimivirus_F (5’-TCGTTTTTACGAAACATGATGG-3’) and MCP_ApolyphagaMimivirus_R (5’-CGATGGTGATTTGGAACACA-3’) (**Arthofer et al 2022**). Cycling conditions were 95°C for 3 minutes; 40 cycles of 95°C for 30 s, 59°C for 30 seconds, and 72°C for 30 seconds; final step at 40°C for 5 minutes. Symbiont 16S rRNA gene copies were quantified using 0.125 µM each of Chl40F/SigF2 (5’-CRGCGTGGATGAGGCAT-3’) and Chl523R (5’-CCYYMCGTATTACCGCAGCT-3’) primers (**Haider et al 2008; Schwarzhans et al 2026**) targeting conserved 16S rRNA gene regions in chlamydial endosymbionts. Cycling conditions were 95°C for 2 minutes; 40 cycles of 95°C for 40 seconds, 64.5°C for 40 seconds, and 72°C for 40 seconds; final step at 40°C for 5 minutes. Each run included a no-template control and positive controls (genomic DNA from *Parachlamydia* PAVD or mimivirus). Genomic DNA was serially diluted in nuclease-free water up to 100-fold and used as template. Reactions (12 µL) contained 4 µL QIAcuity™ EvaGreen 3× dPCR MasterMix (QIAGEN), primers as specified, and 2 µL template DNA. Amplification and detection were performed on a QIAGEN QIAcuity system using the manufacturer’s software.

### Virucidal assays

Viruses used for the virucidal assays were Tromsoevirus (*Marseilleviridae*), megavirus tromsoensis strain A (*Mimiviridae*) (**Queiroz et al 2026**), *Escherichia* phage Polaria (Dutra et al, unpublished) and *Carnobacterium* phage cm44 (Rey et al, unpublished). The virucidal effect was tested by exposing a stock solution of the viruses to 125 µg/ml of Mimicin or of its components MRP-17, SIM-31 and VTG-17 for the cases in which Mimicin was deemed virucidal. Sterile Milli-Q water was used in the controls. To test an immediate antiviral effect, the virus-molecule mixtures were vortexed and serially diluted immediately, while to test the effect of a long incubation, the mixtures were left at room temperature (25ºC) for 30 minutes before being processed. The dilutions were used for titrating the viruses using the TCID_50_ method in *A. terricola* (Tromsoevirus and megavirus tromsoensis strain A) or by double-agar plaque assays in *E. coli* (*Escherichia* phage Polaria) or *C. maltaromaticum* (*Carnobacterium* phage cm44).

### Toxicity assays

Toxicity of all 18 synthesized peptides was tested in the *Artemia salina* model and in anticancer assays using the following cell lines: MRC-5 (lung fibroblasts, ATCC CCL-171™), MCF7 (human breast carcinoma, ATCC HTB-22) and A2058 (human melanoma, ATCC CRL-11147). Artemia tests were made according to **Haug et al 2004** while human cell lines were tested in anti-proliferative assays according to **Schlüter et al 2024**. Cytotoxicity to amoeba trophozoites was qualitatively tested by exposing *A. terricola* (formerly *A. castellanii*) and *A. polyphaga* to 128, 64 and 32 µg/ml of all 18 synthesized molecules in 96 well plates containing 12.000 cells per well. Appearance of cytopathic effects and cell density were verified by observation in an inverted microscope. Cells were grown in PYG medium.

### Sequence analysis and prediction of Mimicin structural features

The Mimicin nucleotide sequence was queried against the NCBI database using BLASTn, employing the megablast algorithm and default parameters. Significant hits were downloaded in FASTA format and aligned using MAFFT (**Katoh and Standley 2013**) with the auto strategy. The same approach was applied to the Mimicin protein sequence, using BLASTp with standard parameters, the protein-protein blast algorithm, and the ClusteredNR (nr_cluster_seq) database. Only BLASTp results with an e-value lower than 10^−3^ and identity over 45% were considered. In cases where multiple hits corresponded to the same species in either BLASTn or BLASTp, the result with the best alignment metrics (based on score, e-value, identity, and gaps) was selected. To carry out the alignments, the sequences obtained exclusively from the BLASTn analysis were translated using the Expasy Translate tool (https://web.expasy.org/translate/) on standard mode. The “5’3’ Frame 1” output was used for subsequent analysis. The search was conducted across different domains of cellular life and among viruses. Alignment visualization was carried out using MEGA 12 software (**Kumar et al 2024**).

Mimicin, MRP-17, SIM-31, and VTG-17 were subjected to structural modelling. All options were inserted into AlphaFold Server, powered by AlphaFold 3 (**Abramson et al. 2024**). Each output was then analyzed in FoldSeek (https://search.foldseek.com/search) (**van Kempen et al. 2023**), considering different databanks such as BFVD, AFDB-proteome, AFDB-swissprot, AFDB50, BFMD, CATH50, GMGCL_ID, MGNIFY_ESM30 e PDB100. Visualization was carried out with PyMOL 3.1 (https://pymol.org/). Homology and structure prediction were also made using the PDB_mmCIF70_20_Feb structural database from HHpred (**Soding et al 2005**). The presence of signal peptides was verified by the SignalP 6.0 server (**Teufel et al. 2022**). The parameters used were the “other” for organism, and the slow mode for the model option. Topology analysis was verified using the DeepTMHMM tool (**Hallgren et al. 2022**). Intrinsic protein disorder was predicted using PONDR (**Xue et al 2010**). AMP features were based on predictions and calculations made at the APD6 antimicrobial peptide database of the University of Nebraska (https://aps.unmc.edu/home, January 2026 update) (**Wang et al 2016**).

### Transcriptome and metagenome analysis

We mapped all predicted ORFs from the Mimivirus genome onto publicly available transcriptome reads from **Zhang et al 2026** (Bioproject: PRJNA1154642) using CoverM v. 0.6.1 with a minimum covered fraction of 50% and a TPM method for coverage calculation (**Aroney et al 2025**). The TPM values from CoverM were converted to log10 with R and plotted using ggplot2. For metagenomic analysis, we used methods similar to those described previously (**Rodrigues et al 2025**). Briefly, predicted proteins from 16,801 metagenomes from the SPIRE database (**Schmidt et al 2024**) were searched against the Mimicin protein using lastal v. 959. Only contigs >5 kbp in length were used for this analysis, and best matches were examined manually.

## Results

### Antimicrobial peptides can be predicted *in silico* from giant virus proteins and are enriched in mimiviruses and marseilleviruses

When all available deposited proteomic information was used as source for predicting AMPs, 155.901 hits were found. These were divided into: 100.538 for Imitervirales, 37.355 for Algavirales and 18.008 for Pimascovirales. Since the high number of putative AMPs found could be related false positives due to the large genomes of giant viruses, we considered different AMPValidate probabilities as cut-offs and corrected the number of potential AMPs with the genome size of viruses that infects amoeba as a mean to normalize the data. Instead of seeing a direct correlation between the number of predicted AMPs to genome size, that could be taken as an indication of purely false positives, it was noted that mimiviruses have a higher density of predicted AMP sequences per genome size than the other groups of viruses of amoeba (**Figure 1**). When no cut-off is applied, the family *Mimiviridae* has the highest average density of 194±44 predicted AMPs/100Kb. The density is also high for mimiviruses with the 50% cut-off (83±22 AMPs/100Kb). The second viral family with a high density of predicted AMPs was *Marseilleviridae* (179±122 with no cut-offs, 89±50 with 50% cut-off). In the case of marseilleviruses, insectomime virus is a clear outlier, resulting in a large standard deviation. However, even if insectomime virus is removed from the analysis, marseilleviruses remains the second group with a high AMP density (146±11 with no cut-offs and 75±8 with 50% cut-off) (data not shown).

**Figure 1:**
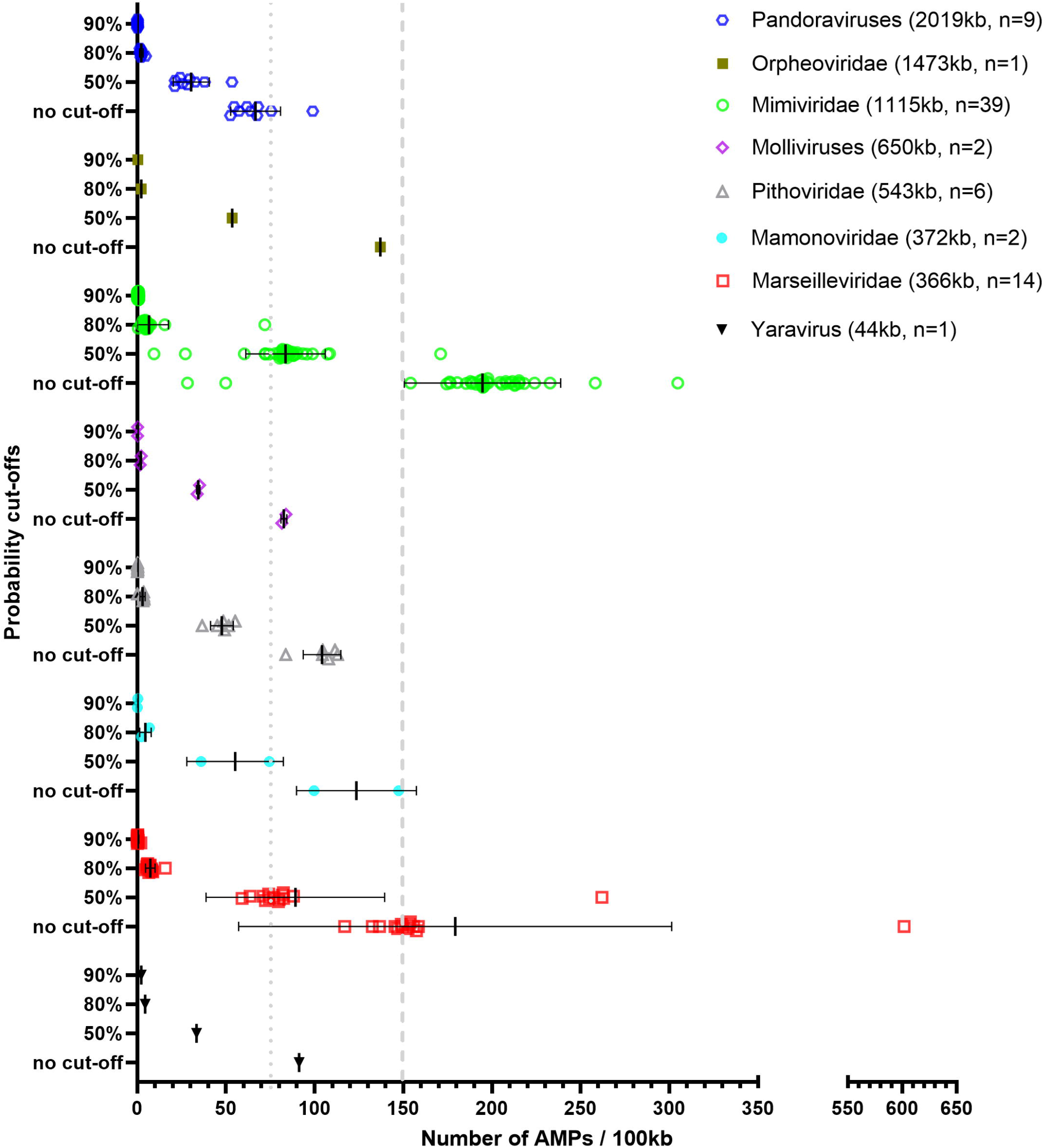
Density of predicted AMPs in different groups of viruses of amoeba. The output of of AMPs predicted by PyAMPA was normalized by genome size and shown as the number of AMPs per 100Kb of genome either as raw predictions (no cut-off applied) or after the application of increasing cut-offs based on AMPValidate probabilities (50, 80 and 90% confidence). The dotted and dashed lines show arbitrary 75 AMPs/100Kb and 150 AMPs/100Kb marks, respectively. Viral groups are plotted in decreasing average genome size from the top to the bottom of the figure. The legend indicates the virus groups in question, the average genome size from the isolates tested and the number (n) of isolates included in the analyses. The mean and standard deviations are indicated in the graph.

### Mimicin, an uncharacterized small protein from mimivirus, has broad antibacterial and differential antiviral activities

One small protein of 74 amino acids (Mimicin) and seventeen predicted AMP sequences, including three internal fragments of Mimicin, were selected for peptide synthesis (**Supplementary Table 1**). When tested in antimicrobial assays, fourteen of the synthesized peptides had no observable effect against the bacteria and yeast strains tested. In contrast, Mimicin had the strongest antimicrobial activity followed by the three predicted AMPs within its sequenced (**Figure 2**). These active AMPs were named MRP-17, SIM-31 and VTG-17 and comprise most of the Mimicin protein (**Supplementary figure 1**). Only Mimicin and SIM-31 had antimicrobial effect against all tested strains, while MRP-17 and VTG-17 had only marginal effects. A compilation of the antimicrobial results is shown in **Table 1**. Neither Mimicin or the seventeen predicted AMPs were toxic in the *Artemia salina* or in a human cancer cell line models at the concentration of 128 µg/ml. Mimicin and SIM-31 were slightly toxic to *A. terricola* and *A. polyphaga* cells, qualitatively affecting part of the cell populations from the 128 to 32 µg/ml doses in the first day following exposure, but with a recovery over time (data not shown).

**Table 1:** Antimicrobial activity expressed as the MIC values of Mimicin, MRP-17, SIM-31 and VTG-17 expressed in µg/mL or in µM. Values indicate the lowest dilution in which complete eradication of the bacteria or yeast tested was verified. Nd: antimicrobial effect not detected.

|  | Species | Mimicin |  | MRP-17 |  | SIM-31 |  | VTG-17 |  |
| --- | --- | --- | --- | --- | --- | --- | --- | --- | --- |
|  |  | µg/mL | Micromolar | µg/mL | Micromolar | µg/mL | Micromolar | µg/mL | Micromolar |
| Gram + | *B. subtilis* | 16 | 1.77 | 32 | 14.96 | 8 | 2.1 | 128 | 66.7 |
|  | *S. aureus* | 32 | 3.54 | nd | nd | 32 | 8.4 | nd | nd |
|  | *S. borealis* | 4 | 0.44 | nd | nd | 8 | 2.1 | 32 | 16.67 |
|  | *S. epidermitis* | 16 | 1.77 | nd | nd | 8 | 2.1 | nd | nd |
| Gram - | *E. coli* | 16 | 1.77 | 128 | 59.85 | 8 | 2.1 | nd | nd |
|  | *K. pneumoniae* | 32 | 3.54 | nd | nd | 32 | 8.4 | nd | nd |
|  | *P. aeruginosa* ATCC | 16 | 1.77 | 128 | 59.85 | 16 | 4.2 | nd | nd |
|  | *P. aeruginosa* PA01 | 16 | 1.77 | 128 | 59.85 | 8 | 2.1 | nd | nd |
| Yeast | *Candida albicans* | 128 | 14.16 | nd | nd | 128 | 33.62 | nd | nd |

**Figure 2:**
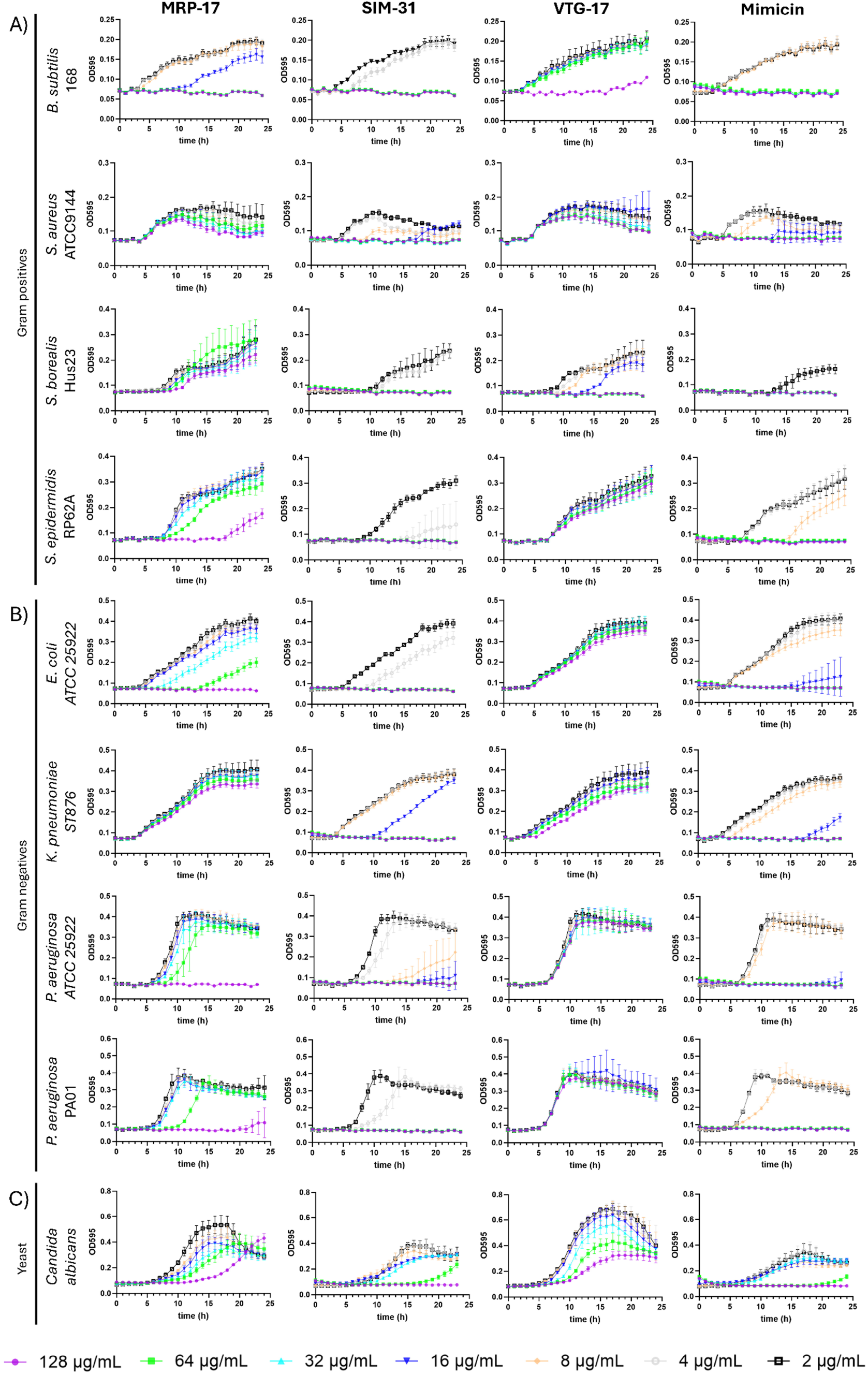
Antimicrobial activity of Mimicin, MRP-17, SIM-31, and VTG-17. Antimicrobial activity was measured by exposing Gram positive bacteria (A), Gram negative bacteria (B) or a yeast (C) to increasing concentrations of each molecule and measuring OD595 of the cultures over 24 hours. All conditions were tested in triplicates and plotted as mean plus standard deviation. Doses lower than 2 µg/ml were omitted from the graphs since the lowest dose observed with antimicrobial effect was 4 µg/ml in few cases.

*B. subtilis* biosensor strains were used to evaluate the antibacterial mechanisms of action of Mimicin, MRP-17, SIM-31 and VTG-17. Both Mimicin and SIM-31 affected bacterial membrane integrity in a concentration-dependent manner, with a more potent effect seen for Mimicin, while MRP-17 and VTG-17 had no effect on the membrane integrity biosensor strain (**Supplementary Figure 2**). In addition, Mimicin and SIM-31 produced a response in the *liaG* viability reporter at concentrations ranging from 25 to 100 µg/mL, indicating an effect on cellular viability. No significant signals were noted in any of the other eight biosensors used when exposed to Mimicin, MRP-17, SIM-31 and VTG-17, indicating that no other bacterial stress response is activated by them in *B. subtilis* the concentrations tested.

The virucidal effect of Mimicin was tested against Tromsoevirus (*Marseilleviridae*), megavirus tromsoensis strain A (*Mimiviridae*), *Escherichia* phage Polaria (*Drexlerviridae*) and *Carnobacterium* phage cm44 (Class *Caudoviricetes*, unassigned family). The phages were used as viruses not related to nucleo-cytoplasmic large DNA viruses. No virucidal effect was noted against either giant virus, even after an incubation of 30 minutes with 125µg/ml of Mimicin (**Figure 3A-B**). In contrast, both phages were affected by Mimicin and SIM-31. *Escherichia* phage Polaria was completely inactivated after incubation for 30 minutes with Mimicin (**Figure 3C**). Because of this finding we also tested MRP-17, SIM-31 and VTG-17, revealing that only the SIM-31 portion of Mimicin has a virucidal effect against *Escherichia* phage Polaria (**Figure 3D**). *Carnobacterium* phage cm44 was partially affected by Mimicin after immediate exposure, and either partially inactivated by SIM-31 or fully inactivated by Mimicin after an exposure of 30 minutes (**Figure 3E**). MRP-17 and VTG-17 had no effect on any of the phages regardless of the incubation time.

**Figure 3:**
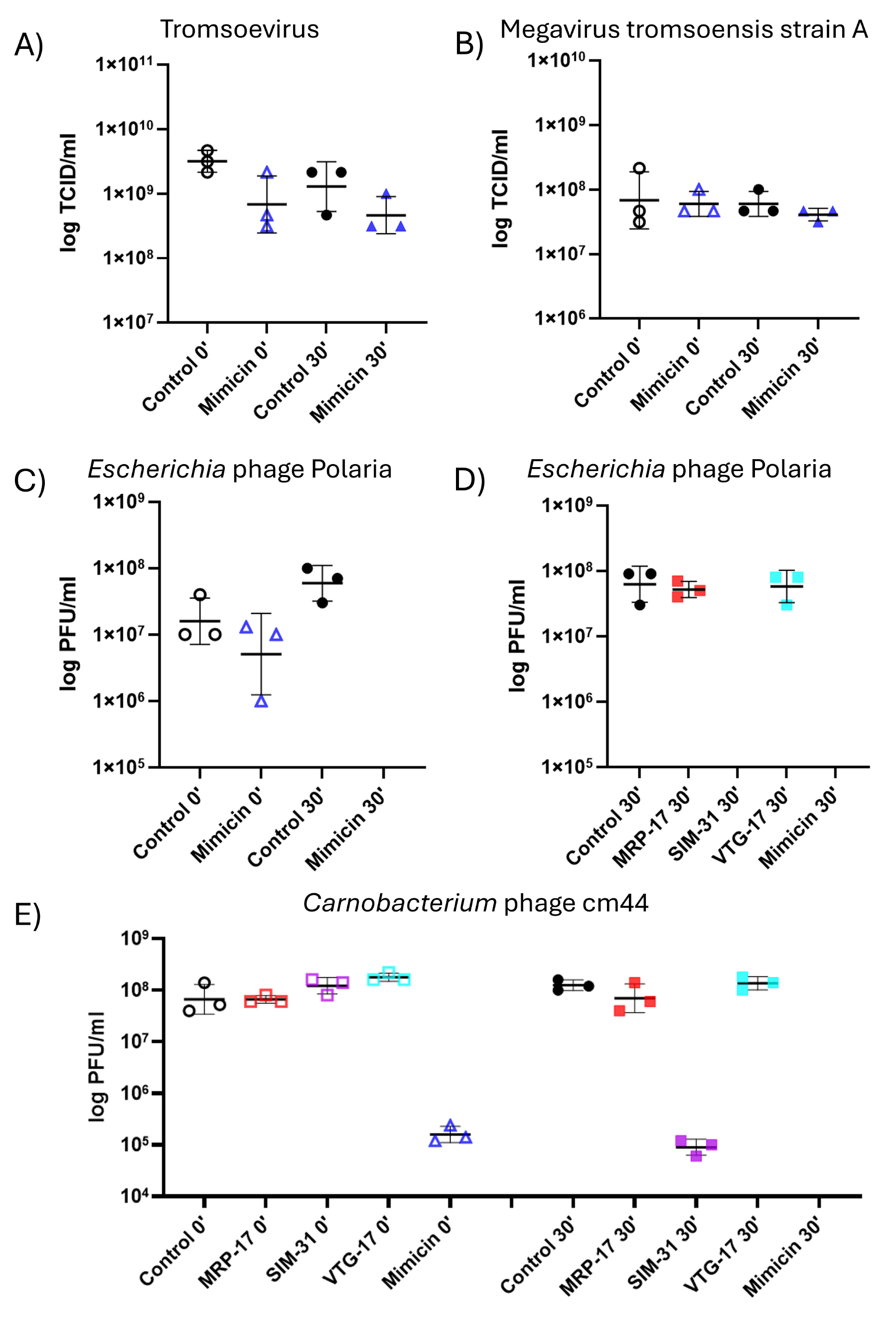
Virucidal activity tests. The virucidal activity of Mimicin, either as immediate exposure or after an incubation of 30 minutes, was tested against Tromsoevirus (A), megavirus tromsoensis strain A (B) and *Escherichia* phage Polaria (C). Additionally, the virucidal effect of MRP-17, SIM-31 and VTG-17 was tested against *Escherichia* phage Polaria after 30 minutes of incubation (D). All four molecules were tested as immediate exposure or after an incubation of 30 minutes against *Carnobacterium* phage cm44 (E). All conditions were tested in triplicates and plotted as individual values with the mean and standard deviation evidenced.

### Mimicin and SIM-31 have species-specific antibacterial activity against chlamydial endosymbionts of amoeba

Before testing the activity of Mimicin and SIM-31 on bacterial endosymbionts of *Acanthamoeba*, a quantitative evaluation of their toxic effect for *A. terricola* trophozoites was carried out in the experimental settings to be used. No significant toxic effect was noted for *A. terricola* cells exposed for 30 minutes to Mimicin or to SIM-31 at 32 µg/ml during a monitoring period of 72 h (**Supplementary Figure 3**). Next, antibacterial activity against bacterial symbionts was tested. Because symbiont relationships vary between different bacterial symbionts (**Collingro et al 2020**), we studied the effect of the AMPs on a highly virulent parasitic symbiont, *Parachlamydia acanthamoebae* (hereafter *Parachlamydia*), and a milder parasite, *Protochlamydia amoebophila* (hereafter *Protochlamydia*), respectively. To this end, we pre-exposed the bacterial symbionts to Mimicin or SIM-31 at 32 µg/ml before infecting *A. terricola*. The molecules were thus able to interact with the symbionts directly before the infection and with the infected amoeba trophozoites for a total of 30 minutes, when the incubation media was replaced. In *Acanthamoeba, Parachlamydia* and *Protochlamydia* establish intracellular inclusions in *Acanthamoeba* and differentiate into reticulate bodies (RBs) within 12-24 and 24-48 hpi, respectively; the first elementary bodies (EBs) emerge at ~24 hpi for *Parachlamydia* and ~72 hpi for *Protochlamydia* (**Greub & Raoult 2002; Horn 2008; König et al 2017**). Our observation period thus encompassed both chlamydial developmental stages.

*Protochlamydia* showed a clear reduction in symbiont numbers compared to the control when treated with Mimicin or SIM-31. The inhibitory effect was strongest at 48 and 72 hours post infection (hpi) (p<0.005; two-way ANOVA) (**Figure 4A**). After entry, an increase in symbiont numbers was observed only in the untreated condition. Under Mimicin exposure, symbiont counts remained essentially constant through 72 hpi, indicating markedly reduced intracellular multiplication. Notably, the low symbiont abundance coincided with improved amoeba growth in treated conditions, with a larger and significant increase in the presence of Mimicin (p<0.0005; two-way ANOVA) than SIM-31 (non-significant) at 48 and 72 h pi (**Figure 4B**).

**Figure 4:**
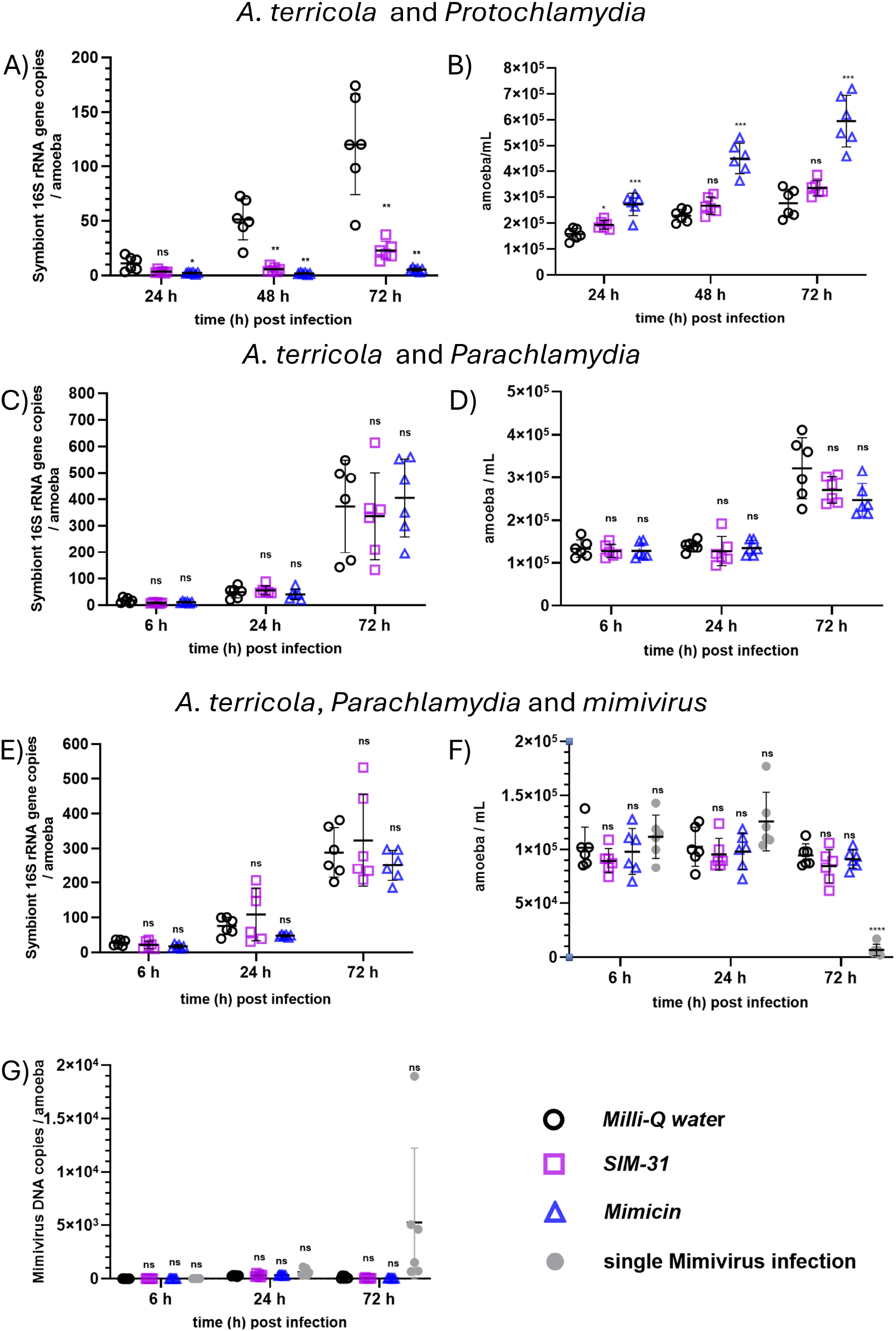
Antibacterial activity of Mimicin and SIM-31 against bacterial endosymbionts of amoebae. Elementary bodies of *Protochlamydia* and *Parachlamydia* symbionts were preincubated with Mimicin or SIM-31 at 20 °C for 30 minutes. *A. terricola* was then infected with the peptide-symbionts mixture. At the indicated times post infection, cells were collected and DNA copies were measured by dPCR. A) *Protochlamydia* genome copies per amoeba. B) *A. terricola* cell counts from co-cultures with *Protochlamydia*. C) *Parachlamydia* genome copies per amoeba. D) *A. terricola* cell counts from co-cultures with *Parachlamydia*. E) *Parachlamydia* genome copies per amoeba during mimivirus-*Parachlamydia* coinfection. F) *A. terricola* cell counts during mimivirus-*Parachlamydia* coinfection. G) Mimivirus genome copies per amoeba cell during mimivirus-*Parachlamydia* coinfection. Statistical significances (*, p<0.05; **, p<0.005; ***, p<0005, ****, p<0.0001) by two-way analysis of variance with Dunnett’s post-hoc test are indicated. Displayed data represents means and standard deviations of two independent experiments (each performed in triplicate).

In contrast, for *Parachlamydia*, a symbiont known to be able to inhibit mimivirus (**Arthofer et al 2022**), no statistically significant change in symbiont numbers and amoeba counts was observed during the infection between treated and untreated samples (**Figure 4C-D**). This indicates a species-specific sensitivity to the tested mimivirus AMPs. We thus tested the effect of the AMPs on amoeba coinfection with *Parachlamydia* and mimivirus. No effect of Mimicin or SIM-31 was seen for *Parachlamydia* or mimivirus loads as well as in amoeba counts, when treated with Mimicin or SIM-31 (**Figure 4E-G**). The only difference was observed between amoeba cell numbers between cultures infected with mimivirus alone and the co-infected cultures (**Figure 4F**), in which the absence of *Parachlamydia* led to increased amoeba cell death at 72 hpi.

### Mimicin is coded by a taxonomically restricted mimivirus gene expressed early during the infection cycle

Mimicin was originally annotated as a hypothetical protein from Amazonia virus, a mimivirus isolated in Brazil (**Assis et al 2015**). BLASTn searches found a few corresponding sequences from other mimiviruses isolates, including the original isolate from France, and matches to large proteins from other organisms (**Figure 5A**). In most mimivirus genomes where this protein sequence was identified, existing annotations either failed to recognize its presence or did not predict it in the correct open reading frame. Among organisms in which the full protein was correctly annotated, especially cellular organisms, functional assignments were inconsistent, with most entries being annotated as hypothetical proteins. Given the limitations that arise from absence from functional information and weak sequence similarity, our current analysis does not provide sufficient phylogenetic resolution to draw reliable conclusions regarding Mimicin’s origins or evolutionary history. Importantly, complete or partial forms of MRP-17, SIM-31 and VTG-17 were detected in all organisms included in the alignment, indicating a broad distribution of these motifs occurring together across the analysed taxa.

**Figure 5:**
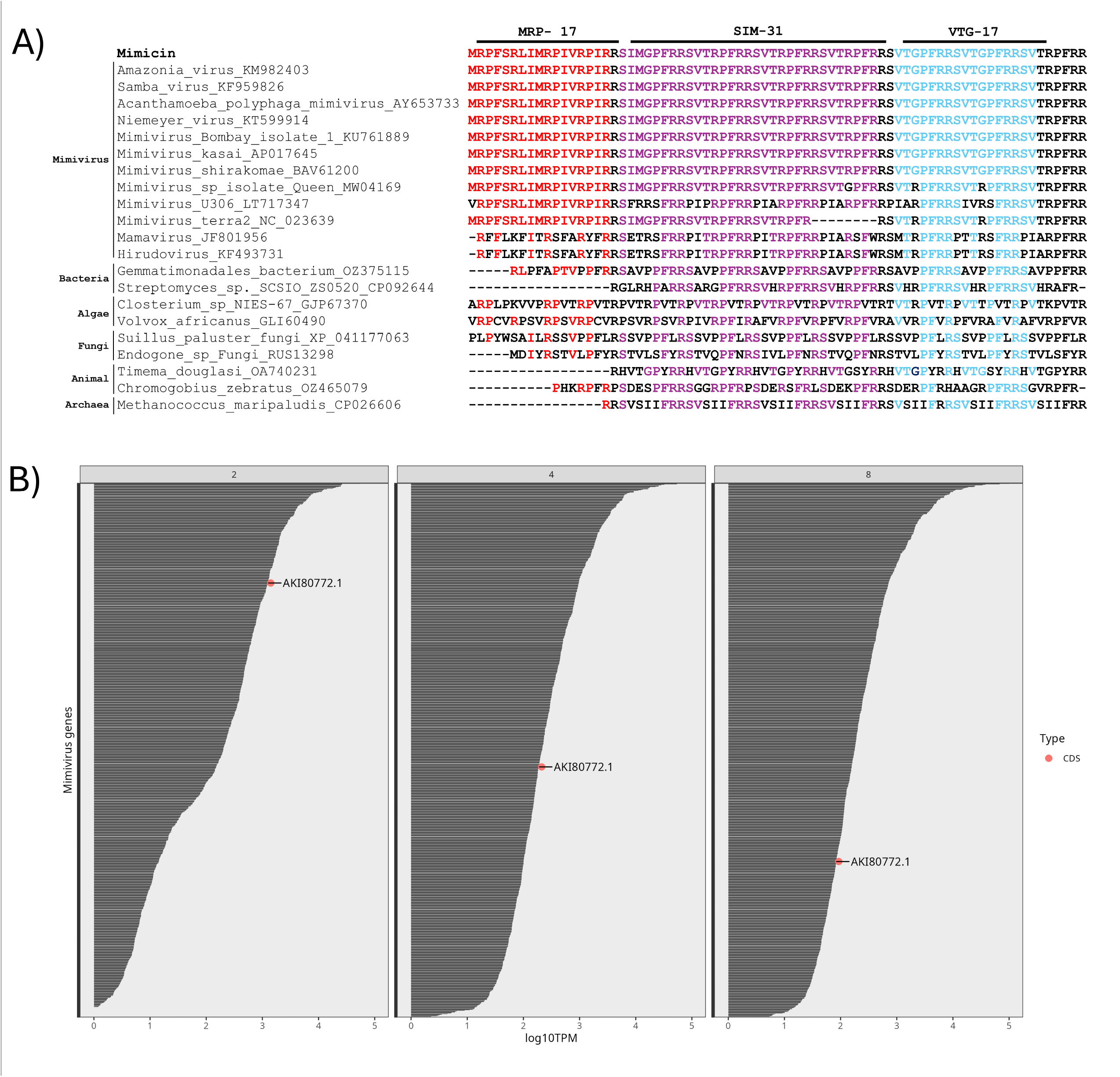
Mimicin sequence diversity and expression. A) Alignment between Mimicin and its BLASTn and BLASTp matches. The regions comprising MRP-17, SIM-31 and VTG-17 are highlighted in red, purple and blue, respectively. Note that excluding mimiviruses, proteins from other organisms are larger and only the aligned regions are shown. Accession numbers of each aligned sequence are shown in the figure. B) Ranking of mimivirus transcripts based on the RNAseq data from **Zhang et al 2026**. Each graph shows a different timepoint after infection (2hpi on the left, 4hpi on the middle and 8hpi on the right), with Mimicin indicated with a red dot.

According to transcriptomic data from **Zhang et al 2026**, Mimicin is an early gene expressed at a relatively high level, especially in the early time-points of infection. When ranked based on abundance of all transcripts, Mimicin shifts from the 184 to the 523 then the 697 position at the 2,4-and 8-hpi timepoints, respectively (**Figure 5B**). A metagenome survey covering >15.000 datasets did not recover any relevant matches to Mimicin. The small size of the protein and its potentially fast evolutionary rate may explain this finding.

### Mimicin do not contain signal peptides, is likely not associated to membranes, and may lack a defined fold

Mimicin lacks cysteine residues and no signal peptide was found *in silico*, indicating that the protein may be active in an intracellular environment. No clear topology features were found *in silico*, indicating no association to membranes and a cytoplasmic location. Mimicin, MRP-17, SIM-31 and VTG-17 have positive net charges (+24, +5, +10 and +4, respectively) and are rich in arginine with the exception of VTG-17. Hydrophobic ratios are 31% for Mimicin, 47% for MRP-17, and 29% for SIM-31 and VTG-17. HHpred found only a low probability (1.5%, e-value of 2900) domain from Mimicin amino acids 2-19 matching penaeidin-3a, a shrimp AMP (**Yang et al 2003**). A single alpha-helix was predicted by AlphaFold3 for Mimicin, while no structure matches were found using FoldSeek. Although some putative structures were predicted by different approaches for the individual peptide regions, the lack of validated homologous targets make these predicted folds speculative. PONDR analysis predicted that Mimicin is a highly disordered protein, with 72 of its 74 amino acids deemed disordered, corroborating the lack of predicted folds for it.

## Discussion

Giant viruses of amoeba are in a constant conflict with other microbes within its host cells (**Karnkowska et al 2026**; **Schulz et al 2025**) and likely evolved strategies to overcome competitors. Here we show that there are many potential AMPs coded by giant viruses, specially within the *Mimiviridae* and *Marseilleviridae* families, where they seem to be enriched. Synthesis of a small number of AMP candidates and subsequent antimicrobial bioassays led to the discovery of Mimicin and three active AMPs within its sequence (MRP-17, SIM-31 and VTG-17). There is no evidence that Mimicin is cleaved into the smaller peptides, and likely it is functional as a whole protein with contributions from each AMP portion, with the second (SIM-31) being the strongest.

Mimicin is coded by an early gene found only in few isolates of mimiviruses, with low similarity to proteins from other taxa including bacteria, archaea, fungi, algae and animals. It is among the non-essential genes lost by the mimivirus M4 strain after repeated passages in axenic conditions made by **Boyer et al 2011**, another indication of it being an antimicrobial molecule. Protein structure prediction analysis suggests that Mimicin is highly disordered, consistent with a lack of fold prediction by Alphafold3 besides a single alpha helix. Intrinsically disordered proteins are common in nature and are often functional, by either folding on binding to their targets or as flexible linkers (**Dyson & Wright 2005; Xue et al 2010**). Intrinsically disordered antimicrobial peptides have already been described from non-viral sources (**Latendorf et al 2019; Cai et al 2022**) and are known to be unstructured in aqueous solutions until conditionally folding close to their ligands (**Bello-Madruga & Burgas 2024**). Intrinsically disordered proteins are also important in the context of giant virus infections, functioning as drivers of phase separation and scaffolding proteins of viral factories (**Rigou et al 2025**).

Mimicin and its components (MRP-17, SIM-31 and VTG-17) are not toxic to human cells and were shown to be antimicrobial *in vitro* against different bacterial strains and against one yeast strain, with a stronger activity noted for Mimicin and its SIM-31 portion. The most probable mechanism of action for both is fast activity directed against microbial cell membranes. Mimicin and SIM-31 were also shown to possess a differential virucidal activity, not affecting a megavirus and a marseillevirus but inhibiting replication of two different phage isolates. Most antiviral peptides target enveloped viruses, but there are examples of peptides with virucidal activity against non-enveloped viruses as well. In those cases, cationic peptides act by disrupting or aggregating negative charged viral particles (**Neghabi Hajigha et al 2024**). Mimicin and SIM-31 are rich in arginine and have positive net charges, fitting into this model against the non-enveloped tailed phages tested. If this is correct, Mimicin could also function against virophages, serving as a tool for intra-virus competition and potentially adding one more layer of complexity to its activities in nature. One reason for the lack of virucidal effect on a representative of the *Marseilleviridae* and one of the *Mimiviridae* family might be due to the higher complexity of their particles, composed of more than one capsid layer and of fibrils in case of the mimivirus, that could serve as a protection against the AMPs. Additionally, Mimicin and SIM-31 might have affected or masked phage ligands, blocking its recognition of the bacterial host cell, something that is not relevant for giant viruses that employs phagocytosis for viral entry.

In co-cultures of *Acanthamoeba* and bacterial endosymbionts, Mimicin and SIM-31 are active against a common symbiont of amoebae, the mild parasite *Protochlamydia*, with corresponding benefits for the amoeba host growth. It remains unclear, however, whether the peptides act only on the extracellular EBs and reduce their infectivity, or whether they can also pass the amoeba cell membrane and the inclusion membrane to reach the intracellular replicative RB stage of the symbionts. The constant number of *Protochlamydia* from 24 to 72 hpi in the Mimicin treated condition, relative to the increase in numbers in the untreated control, suggests peptide entry and/or sustained intracellular antibacterial activity. Nonetheless, an exclusive effect on EB infectivity that limits subsequent RB expansion cannot be fully excluded. Yet, the peptides did not affect the related *Parachlamydia* symbiont and did not influence their inhibitory effect on mimivirus replication during co-infection of *Acanthamoeba*. In fact, resistance of *Parachlamydia* against the tested peptides might reflect the reported antiviral activity of this endosymbiont against diverse giant viruses **(Arthofer et al., 2022**), while *Protochlamydia* does not appear to trigger an antiviral response against giant viruses (*Marseillevirus viennavirus*, family *Marseilleviridae*) (unpublished data).

In conclusion, here we describe the discovery of Mimicin, a small and taxonomically restricted protein from a mimivirus shown to possess antimicrobial activity *in vitro* and against a bacterial endosymbiont of amoeba in co-culture settings. Mimicin and its components are only four of many potential AMPs predicted from giant viruses’ proteins, making them the first step into the investigation of giant virus derived AMPs and their relevance for microbial ecology and biotechnology.

## Acknowledgements

We thank Dr. Bernard La Scola (Aix-Marseille University, France) for kindly donating the *A. castellanii* and *A. polyphaga* strains used in Norway, Dr. Jorunn Pauline Cavanagh (Department of Clinical Medicine, UiT, Norway) for kindly donating the *S. borealis* Hus23 strain, Clara de Castro Ribeiro Jardim for technical support in the virucidal assays, Hege Devold for the toxicity tests in *Artemia salina*, and Ida Kristine Østnes Hansen for technical assistance in MS analyses. We gratefully acknowledge the research group Marbio (UiT, Tromsø) and in particular Kirsti Helland for performing the anti-proliferative assays.

This work was funded by the Centre for New Antibacterial Strategies (CANS) of the Arctic University of Norway (Tromsø Forskningsstiftelse ID:18_CANS_AS 311192/A65276). M.H. acknowledges support from the FWF (https://doi.org/10.55776/P37198) and the FWF Cluster of Excellence COE7 (https://doi.org/10.55776/COE7). J.S.A. and A.K.N.A. acknowledge CNPq, CAPES, and FAPEMIG for their financial support

Part of the bioinformatic analysis were performed on resources provided by Sigma2 - the National Infrastructure for High-Performance Computing and Data Storage in Norway. This research was registered in the Brazilian National System for the Management of Genetic Heritage and Associated Traditional Knowledge—SISGEN A581C34.

