## Supplementary figures for "Mimicin, an antimicrobial protein encoded by mimivirus"

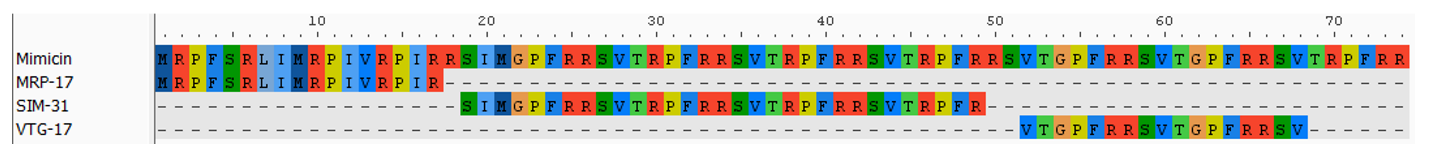

**Supplementary figure 1**: Alignment of Mimicin, MRP-17, SIM-31, and VTG-17 showing the organization of the full Mimicin protein with the three peptides within its sequence.

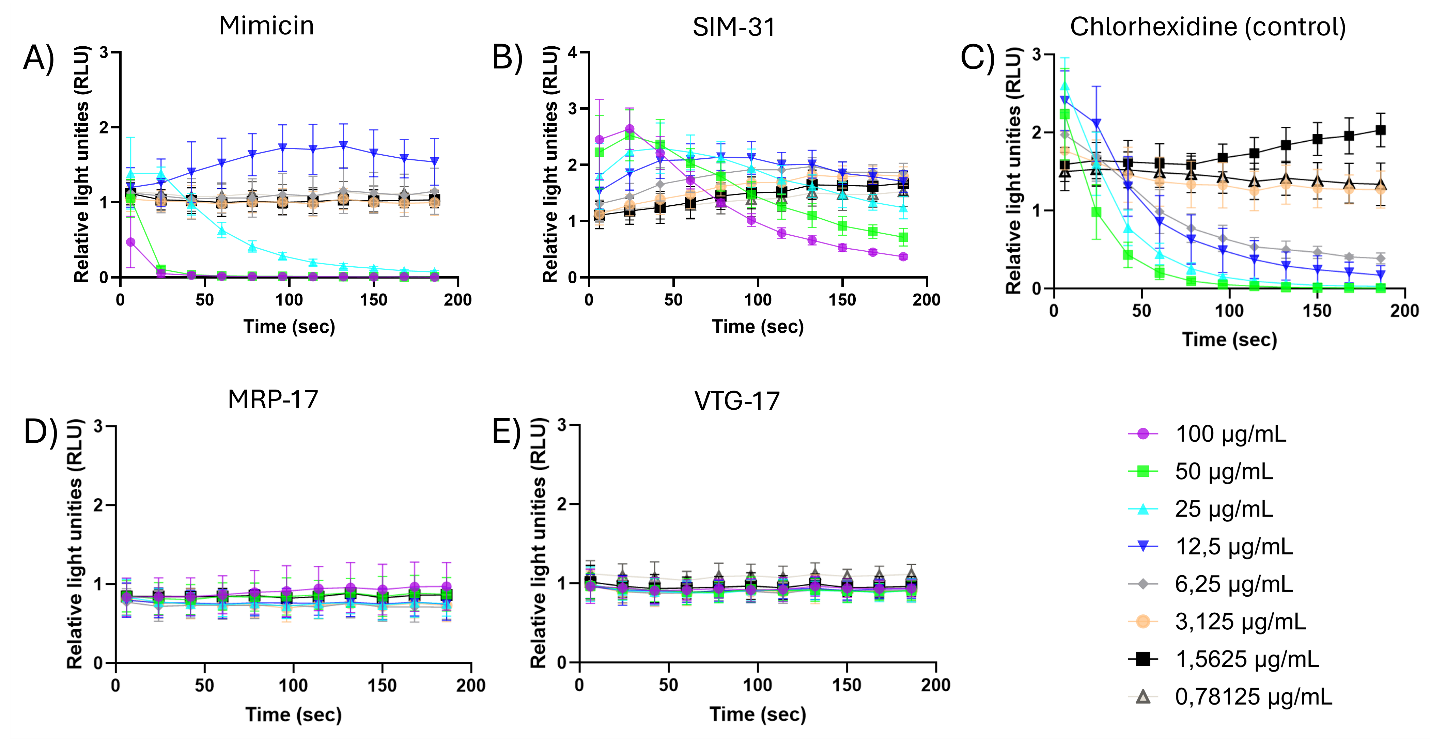

**Supplementary figure 2**: Mechanism of action investigation. Membrane activity of Mimicin (A), SIM-31 (B), chlorhexidine (positive control) (C), MRP-17 (D) and VTG-17 (E). Mimicin led to quick luminescence decline to near-background levels within the first 25–50 seconds at the 100 and 50 µg/mL doses. This immediate collapse of the luminescence signal is consistent with membrane permeabilization and ATP depletion occurring faster than the temporal resolution of the instrument, as commonly observed for highly potent membrane-active compounds. For SIM-31, a pronounced transient luminescence peak was detected within the first ~20 seconds at the 100 µg/mL, followed by a progressive decline to near-background levels by approximately 200 seconds. A similar but less pronounced response was observed at 50 µg/mL. In contrast, MRP-17 and VTG-17 did not produce a concentration-dependent luminescence response, with all tested concentrations indicating little or no detectable membrane-disruptive activity under the conditions tested. All conditions were tested in triplicates and plotted as mean plus standard deviation.

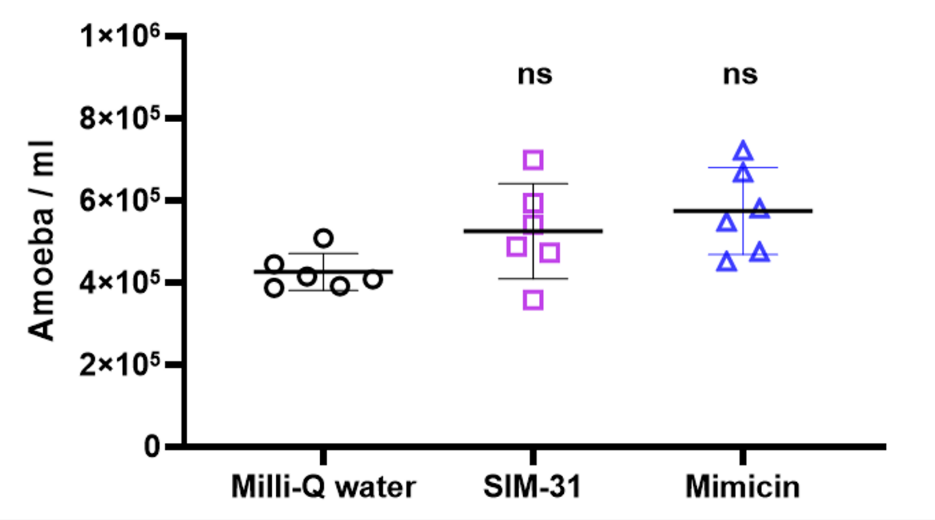

**Supplementary Figure 3:** Effects of peptide preincubation on *A. terricola* in the co-culture experimental setting. The toxicity of peptides to *A. terricola* under preincubation conditions. Amoebae were exposed to peptides for 30 min, then peptides were removed and amoeba numbers were measured using a LUNA cell counter at 72 h. All conditions were tested in six replicates and plotted as individual values with the mean and standard deviation evidenced.
